# *Arabidopsis thaliana* induces multigenerational stress tolerance against biotic and abiotic stressors and memorization of host colonization in *Bacillus subtilis*

**DOI:** 10.1101/2022.05.29.493878

**Authors:** Omri Gilhar, Tsviya Olender, Asaph Aharoni, Jonathan Friedman, Ilana Kolodkin-Gal

## Abstract

*Bacillus subtilis* is a beneficial bacterium that supports plant growth and protects it from bacterial, fungal, and viral infections. Here using a simplified system of *B. subtilis*, and *Arabidopsis thaliana* interactions, we found that history-dependent behavior is a potentially important manifestation of host colonization, worth classifying and quantifying. To study history-dependent adaptation to plant hosts, we develop a simple framework for measuring the physiological memory of *B. subtilis* following its interaction with *Arabidopsis thaliana*. We found that *A. thaliana* secretions reduce the lag time in pre-exposed bacteria compared with naïve *B. subtilis* cells, even after their complete removal. Pre-exposed *B. subtilis* cells colonized plant roots more efficiently than naïve bacteria, and were more resistant to biotic and abiotic stressors such as salicylic acid, and high salinity. Descendants of bacteria treated with plant secretions had an advantage in the competition against unexposed bacteria for root colonization. The effect of plant secretions was independent of their roles as nitrogen and carbon sources. Transcriptome analysis of both ancestors and descendants revealed that a specific set of plant-induced processes, among them c-di-AMP homeostasis, and the general stress response, maintain the signature of association with the plant in descendants of pre-exposed bacteria. Consistently, plant secretions compensated for the loss of c-di-AMP cyclases but required the general stress response and the master regulator Spo0A to exert their short and long-term effects.

Overall, our work demonstrates that bacterial memory manifested by multigenerational reversible adaptation to plant hosts confirms an advantage to symbiotic bacteria during competition.

## Introduction

*Bacillus subtilis* and related *Bacillus* species, such as *B. amyloliquefaciens, B. velezensis and B. mojavensis*, serve as biocontrol agents, plant and mammalian probiotics^1–3^. Beneficial *Bacillus* species protect their hosts from both fungal and bacterial pathogens ^4–6^, a protection that is mediated, in part, by the formation of biofilms ^7,8^, as well as by the production of a wide range of antibiotics ^2,9^. *B. subtilis* also elicits induced systemic resistance (ISR) in plant hosts via its secondary metabolites^10–12^. The successful colonization of plant hosts depends on the capacity of *B. subtilis* to form biofilms and occurs at preferred sites of root exudation, where plants release C- and N-containing compounds into the surrounding soil ^13^. Plant metabolites were shown to induce motility, chemotaxis, matrix production and antibiotic production ^11,14–17^. The plant can also induce long-term adaptations: Bacteria that were repeatedly cultured with *A. thaliana* evolved rapidly, to generate morphotypes with improved root colonization. In this experimental system, *Bacillus subtilis* cells diversified into three distinct morphotypes altered in their growth and pellicle formation in a medium supplemented with plant polysaccharides ^18^.

Plant polysaccharides serve as a signal for biofilm formation transduced by activating kinases controlling the phosphorylation state of the master regulator Spo0A^15^, which directly or indirectly controls the expression of some 500 genes in *B. subtilis*^19,20^. In part, Spo0A regulates the generation of spores, a dormant and resilient cell type. However, sporulation is considered a last resort to cope with extreme environments, as well as biotic and abiotic stressors^21^.

One effective strategy for tackling stressors can be the utilization of information collected during a previous interaction with similar stress. Although it was shown that bacterial behavior could be modified to reflect preceding events, the time elapsed between the events usually is very short, counting in seconds or minutes, and, as such, may not be classified as a “true memory”^22^. Mechanistically, multigenerational responses operate through the transmission of cytoplasmic proteins with lifetimes more extended than the generation of a bacterial cell. For example, the stability of proteins encoded by the lactose operon in *E. coli* reduced the bacterial lag phase during growth in media in which the concentrations of glucose and lactose fluctuated^23^. In addition, a recent study in P. *aeruginosa* reported multigenerational memory based on nucleotide signaling related to abiotic surfaces. In this study, the bacteria formed a cyclic-AMP–TFP-based memory of surface attachment that enabled them and their descendants an improved re-attachment^24^.

Plants (representing biotic hosts and their surfaces) were reported to induce long-term adaptations in bacteria through a genetic selection of distinct morphotypes: *B. subtilis* repeatedly cultured with A. thaliana evolved rapidly to generate morphotypes with improved root colonization. In this experimental system, the bacterial cells diversified into three distinct morphotypes altered in their growth and pellicle formation in a medium supplemented with plant polysaccharides^25^ (Blake et al., 2021). Whether symbiosis or interaction with biotic surfaces induces more transient multigenerational responses and whether these long-term adaptations provide quantifiable fitness advantage remain unknown.

Given the potential ubiquity and significance of long-term adaptation to the host, quantifying the long-term effects of *B. subtilis* interactions with the host could be an important piece of the puzzle of bacterial regulation, survival strategy, and evolution. To this end, we developed a simple framework for measuring the persistence of memory in ancestors and descendants that were exposed to plant secretions, and the amount of information that was maintained in their transcriptome regarding this interaction. Relying on it, we unraveled several new principles shaping long–term effects of the plant on *B. subtilis* beneficial bacterial communities. First, we found that symbiosis is manifested by a long-term transient adaptation of root-associated bacteria to their host, which is not due to host-driven genetic selection. These long-term adaptations decrease the duration of the lag phase for exposed bacteria and their descendants and promote more efficient root colonization. In turn, the optimization of plant colonization promotes the plant growth mediated by the exposed bacteria, compared with naïve bacteria that never interacted with a host. These symbiotic features were pronounced under abiotic stress. Second, we determined that while a global response to the plant involves 25% of the transcriptome, a much smaller core of genes, primarily associated with the general stress response and antibiotic production maintain their expression pattern in free-living descendants of plant-associated bacteria. Finally, we determined that the plant signaling acts upstream of the nucleotide signaling of c-di-AMP.

Our results indicate that there might be a strong fitness incentive toward memory in *B. subtilis*. If cells could use a memory of their previous interactions with the plant to ‘predict’ future interactions with plant hosts, they might improve their odds for long-term survival in the natural habitat of *B. subtilis*, the rhizosphere.

## Results

### Treatment with *A. thaliana* secretions shortens the lag phase of bacteria and their descendants

To assess whether the effect of the plant on bacterial growth is maintained after detachment of plant colonizers from the plant, we monitored the growth of bacteria that were cultured with or without plant secretions in a fresh medium (Fig. 1A). As root secretions were removed prior to culturing, this system was designed to measure short and long-term adaptation to the plant. We found that pre-treatment with plant secretions affected bacterial growth, and specifically, the lag phase was shortened by previous interactions with plant secretions (Fig. 1B). However, the growth rate and the maximal OD of the bacteria were not significantly altered by previous exposure to plant secretions (Fig. S1). Neither pre-treatment with a carbon source (e.g: glucose of sucrose), nor with a nitrogen source (e.g: glutamate and ammonium chloride) affected the lag phage comparably to plant secretions (Fig. S2)

**Figure 1.**
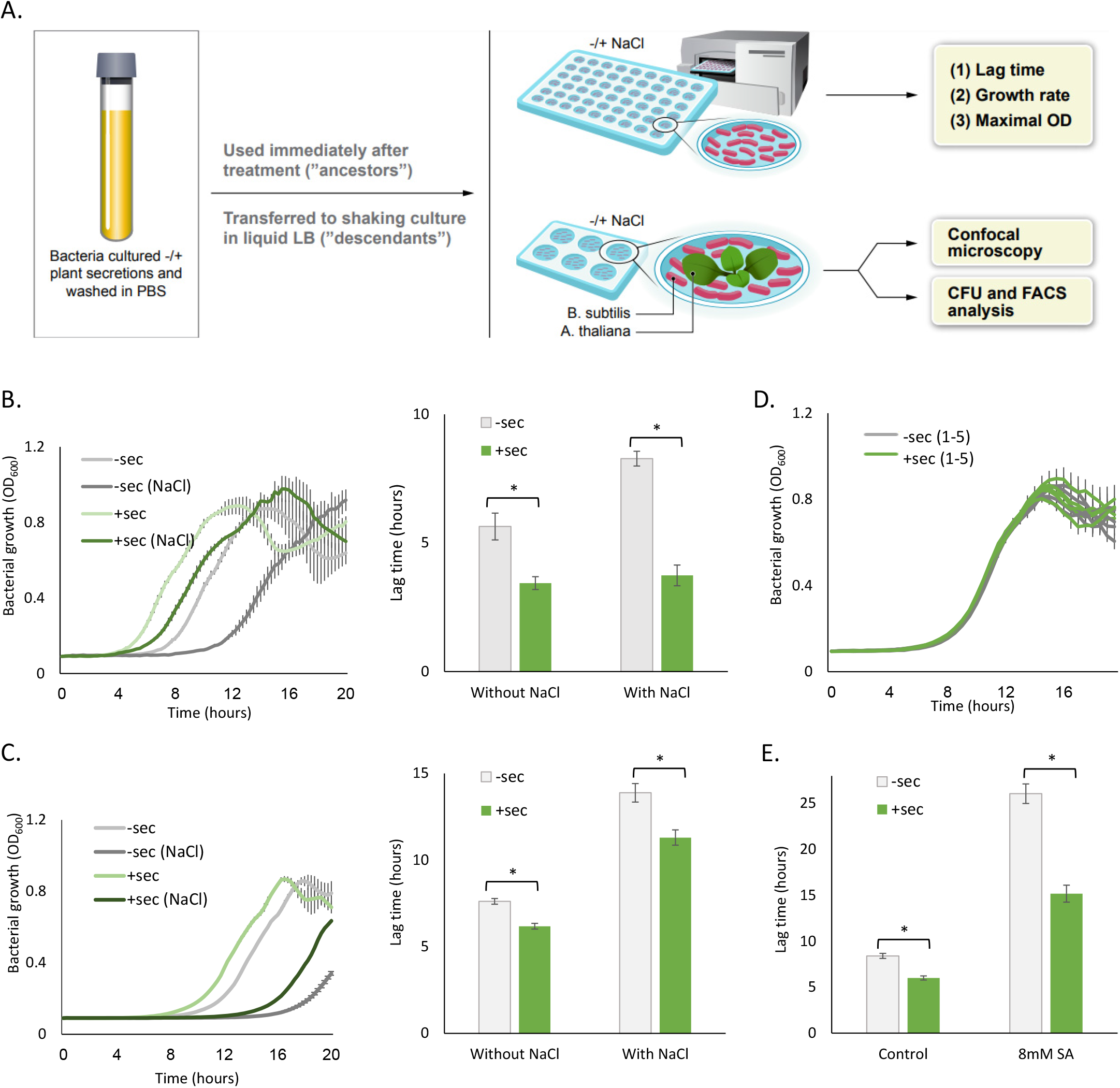
A. *thaliana* secretions grant *B. subtilis* cells increased tolerance to stressors. (A) *B. subtilis* bacteria were cultured in liquid MSgg (“untreated”) or in *A. thaliana* secretions and then washed in PBS. These bacteria were termed “ancestors”. Some of the bacteria were transferred to a shaking culture in liquid LB and grown for four hours. These bacteria were termed “descendants”. The bacteria were then regrown in MSgg and monitored for planktonic growth -/+ 600 mM NaCl in a plate reader. Alternatively ancestors and descendants were cultured with *A. thaliana* plants -/ + 600 mM NaCl, after which root colonization was examined with a confocal microscope, CFU counts and FACS analysis. (B-C) Growth curves and lag time of secretion treated and untreated bacteria (+sec and -sec respectively) and their descendants in the absence and presence of NaCl. Bacteria were grown in a microplate reader as described above and their lag time was calculated (*see materials and methods*) (D) Lag time of bacteria with and without plant secretions in the absence and presence of salicylic acid in “ancestor” bacteria. (E) Growth curves of bacteria that were treated or untreated with secretions, re-streaked separately on LB plates and grown overnight. Single colonies were picked up, transferred to liquid shaking culture for 4 hours at 37°C and then grown in a microplate reader as described above. All experiments were performed in triplicates at least three separate and independent times. Data is expressed as average values ± standard deviations of the means from a representative experiment, performed with replicates. Data was analyzed by two tailed student’s t-test (alpha = 0.05).

To test whether plant secretions promote growth under abiotic stress, we examined whether a more pronounced decrease in lag time is observed under high salinity, one of the most common stressors in the soil and rhizosphere ^26^. The decrease in lag time upon secretion treatment was larger in the presence of NaCl than in its absence (29 percent difference) (Fig.1B). The beneficial effect of pre-exposure to the plant growth was multigenerational-descendants of bacteria treated with secretions grown for 8 generations in rich media exhibited a significantly shortened lag phase compared with descendants of unexposed bacteria (Fig. 1C). However, growing the bacteria for additional 16 generations resulted in the loss of the shortened lag phase in the descendants of the secretion-treated bacteria (Fig. S3). Here again, neither carbon nor nitrogen was sufficient to mimic the beneficial effects of plant secretions to the descendants (Fig. S4).

We then asked whether enhanced growth and stress resistance result from a plant-induced selection. The descendants of secretion treated and untreated bacteria were isolated from single colonies on LB agar, and single isolates were compared for growth. Re-isolated cells from treated and non-treated cultures exhibited a similar growth and a comparable lag time (Figs 1D and S5) to their parental strain. Therefore, the decreased lag time of the bacteria in this system most likely reflects a temporary adaptation to the secretions and does not result from a hereditary change.

To determine whether common stressors other than NaCl induce a more pronounced lag time decrease in secretion-treated bacteria as compared with untreated bacteria, we used high concentrations of salicylic acid, a phytohormone playing an important role in plant protection. The decrease in lag time upon secretion treatment was larger in the presence of salicylic acid than in its absence (32 percent difference) (Fig. 1E).

### Memorization of root secretions increases the competitiveness of *B. subtilis* during root colonization

In the rhizosphere, detachment from the plant is quite common, and the detached bacteria often compete with other rhizosphere free-living members for colonization of an alternate host^27^. We therefore examined the effect of pre-treatment with secretions on root colonization. The bacteria treated with secretions colonized the root more efficiently than untreated bacteria (Fig. 2A). Furthermore, the descendants of secretion-treated bacteria more efficiently colonized the root in high salinity (Fig. 2B).

**Figure 2.**
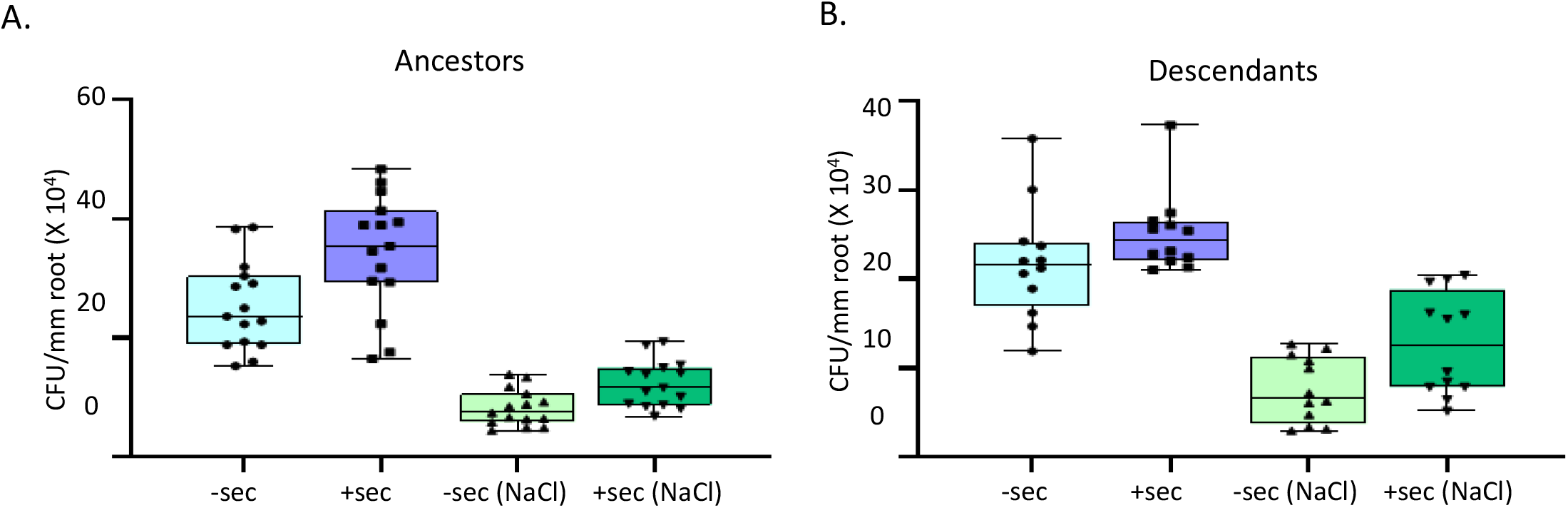
Multigenerational adaptation of *B. subtilis* to *A. thaliana* enhances root colonization in the presence and absence of abiotic stressor. *B. subtilis* colonization of *A. thaliana* and tomato roots after 24h with and without secretion treatment (A-B). Secretion-treated and untreated WT bacteria (+sec and -sec respectively) and their descendants were cultured separately in liquid MSgg with *A. thaliana* seedlings in the absence or presence of NaCl. After 24h, the bacteria colonizing the roots were quantified with CFU counts. Data is expressed as average values ± standard deviations of the means from all independent experiments.

To test whether plant secretions grant the bacteria an advantage during competition with naïve bacteria, we competed naïve bacteria with bacteria exposed to root secretions for root colonization. Fluorescently labeled bacteria were cultured in the absence (mKate-labeled) or presence of plant secretions (GFP-labeled) and then co-inoculated with *A. thaliana* seedlings. As shown, the bacteria pre-treated with secretions colonized the root in significantly more efficiently than untreated bacteria in the absence and presence of NaCl (Figs 2A-C). The bias was not due to the different fluorophores used for tagging as naïve bacteria labeled with GFP and mKate colonized the plant similarly (Fig. S6). Pre-treatment with *A. thaliana* secretions also induced a more efficient colonization of tomato seedlings similarly to *A. thaliana* (Figs 2D and S7). These results indicate that association with a plant allows exposed bacteria and their descendants to colonize a new plant host more efficiently, which differ from the original host that produced the secretions. This effect was also multigenerational as descendants of the treated and untreated bacteria exhibited a similar ratio of root colonization as their ancestors (Fig. 2E).

### Transcriptome profiling reveals a specific set of genes reflecting the multigenerational adaptation of *B. subtilis* to its host

To assess the molecular features of temporal and multigenerational adaptation to the root, we compared the transcriptome of untreated *B. subtilis* cells and cells cultured in plant secretions. *B. subtilis* cells were also cultured in the presence of plant, to examine whether physical association with the root further alters gene expression.

The transcriptome of secretion treatment was similar to bacteria physically associated with the root, indicating that the majority of host-driven gene regulation is due to the exchange of secreted compounds. The number of genes differentially expressed following secretion treatment versus untreated bacteria was 1292, while 1578 genes were differentially expressed following treatment with the plant versus untreated bacteria (Fig. 3A-D, Fig S6). Although the transcriptome architecture of bacteria that physically interact with the plant and bacteria treated with plant secretions alone were similar, with a significant overlap (p < 1.82E-135, Fig. 3B), there was still a differential expression of 508 genes related to processes such as carbon and amino acid homeostasis, c-di-AMP signaling and stress response (Fig. S7).

**Figure 3.**
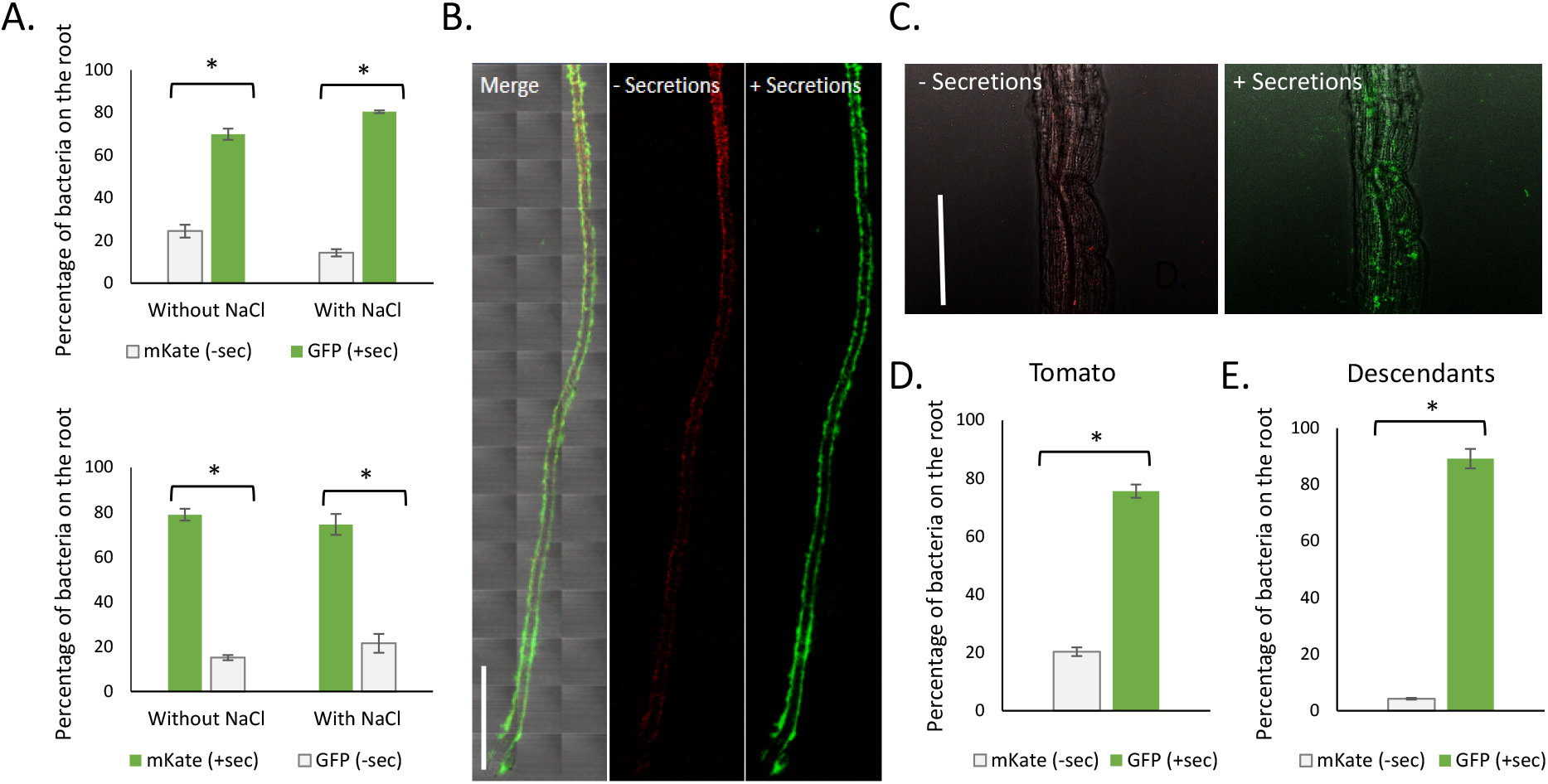
Multigenerational adaptation of *B. subtilis* to *A. thaliana* increases the competitiveness of exposed bacteria and their decedents. (A) Fluorescently-labeled bacteria treated or untreated with secretions were cultured together in liquid MSgg with *A. thaliana* seedlings in the absence or presence of NaCl. The percentage of secretion-treated and untreated ancestor bacteria on the root was quantified with FACS. Representative bright field and fluorescent images of secretion-treated and untreated ancestor bacteria on *A. thaliana* roots in the absence (B) and presence (C) of NaCl. (D) The percentage of secretion-treated versus untreated bacteria on tomato roots without NaCl quantified with FACS (as described above). (E) The percentage of secretion-treated versus untreated descendant bacteria on *A. thaliana* roots without NaCl. Descendant bacteria were grown and treated as described in Fig. 1 and quantified with FACS (as described above). All experiments were performed in triplicates at least three separate and independent times. Data is expressed as average values ± standard deviations of the means from representative experiment, done in technical triplicates. Data was analyzed by two tailed student’s t-test (alpha = 0.05).

Pathway analysis indicated that the physical association with the root as well as exposure to secretions altered the expression of genes related to bacterial life style such as motility and antibiotic production (chance for false analysis= 2.94E-06 and 2.31E-03 respectively) (Fig. 3E). In addition, amino acid homeostasis and iron homeostasis were altered (chance for false analysis = 5.80E-04 and 0.03 respectively). These metabolic alterations could indicate co-feeding relationships between *A. thaliana* and *B. subtilis*. Lastly, we examined the differential expression of genes related to c-di-AMP homeostasis, stress tolerance, biofilm formation and carbon metabolism. Indeed, we observed that interaction with plant secretions induced predominantly upregulation of genes involved in biofilm formation (23 upregulated and 7 downregulated out of 60 genes), stress response (70 upregulated and 13 downregulated out of 200 genes) and carbon metabolism (51 upregulated and 16 downregulated out of 216 genes). Genes affected by c-di-AMP signaling were both up and down regulated (62 upregulated and 60 downregulated out of 443) (Fig. 3F). Notably, our previous results using flow cytometry confirm the induction of the biosynthetic clusters for the antibiotics surfactin and bacillaene by the root, as well as the repression of lactate dehydrogenase^28^.

The alteration of the transcriptome architecture upon exposure to the secretions was widespread. However, the changes in the transcriptome of descendants of pre-exposed bacteria compared with the transcriptome of non-treated bacteria were much more specific (Fig. 3F). Plant-driven induction of metabolic, iron homeostasis biofilm and motility genes were not conserved in descendants (Figs 3E and F). In contrast, host-driven induction of 278 genes was clearly observed in the descendants of treated bacteria (Fig. 3F). These were comprised mostly of genes involved in stress response (89 upregulated) and genes affected by c-di-AMP signaling (15 upregulated and 21 downregulated). Interestingly, 18 genes involved in motility were downregulated in descendant bacteria, unlike ancestor bacteria in which the motility genes were predominantly upregulated (21 upregulated and 5 downregulated out of 42 genes). Overall, these results indicate that specific genes related to stress tolerance are induced in plant-associated bacteria, as well as their descendants (Fig. 3F), and could account for both increased colonization of a new host and resistance to abiotic stress. Furthermore, our results indicate that a specific transcriptional signature reflecting a multigenerational effect of the plant host, and that this response includes relaxed response of the stress response clusters in the descendants of the pre-exposed bacteria compared with their ancestor

### Microbial mutigenerational response involved cyclic di-AMP signaling, the general stress response and the master regulator Spo0A

Cyclic di-AMP is a signaling dinucleotide used for bacterial intracellular signaling. In Gram-positive bacteria, it affects growth, sporulation, virulence, DNA damage-repair, osmoregulation ^29^ and biofilm formation^30^. Di-Adenylate cyclases (DACs) convert pairs of ATP molecules into c□di□MP. In parallel, enzymes degrading c□di□MP play a role in the homeostasis of this second messenger. These two events lead to changes in the c□di□MP cellular pool, which in turn modifies the activities of target proteins ^31–33^. In *B. subtilis*, the enzymes that produce c-di-AMP are designated DisA, CdaA and CdaS, with DisA being a cardinal sensor of DNA damage to activate the general stress response^33,34^. CdaA has a constitutive expression and is localized to the cell membrane, and its paralog CdaS synthesizes c-di-AMP during spore formation, expressed only during sporulation and is localized to the forespore^35^. *B. subtilis* also secretes c-di-AMP, which can act as an extracellular signal to affect biofilm formation and plant attachment^30^. Whether symbiosis also induces more transient multigenerational responses, and whether c-di-AMP plays a role in such long-term adaptations remains unknown. However, c-AMP was shown to mediate multigenerational memory of artificial surfaces in the pathogen P. *aeruginosa*, linking nucleotide signaling with microbial memory^24^.

We therefore tested the role of the diadenylate cyclases in the adaptation of *B. subtilis* to the plant. The parental strain and its diadenylate cyclase mutants (*ΔdisA, ΔcdaA* and *ΔcdaS)* were cultured with and without secretions. Interestingly, *ΔcdaA* exhibited a significantly longer lag time and a decreased root colonization as compared to the WT in the absence of secretions (*p* < *0.05*), but following treatment with secretions, the lag time (Fig. 5A) was comparable to the parental strain (*p* = *0.25*). The lag time decrease of *ΔcdaS* following secretion treatment was not significantly different from the WT in ancestors (*p* = 0.75) and descendants (*p* = 0.53) (Figs. 5A and B). This result indicates that root secretions compensate for the growth defects in the diadenylate cyclase mutant *ΔcdaA*. Consistent with the multigenerational effect on the transcription of genes related to c-di-AMP homeostasis, the effect was conserved in descendants of pre-treated bacteria (Fig. 5B). These results indicate that the plant secretions can compensate over the loss of diadenylate cyclase in the bacteria exposed to secretions, as well as in their descendants.

**Figure 4.**
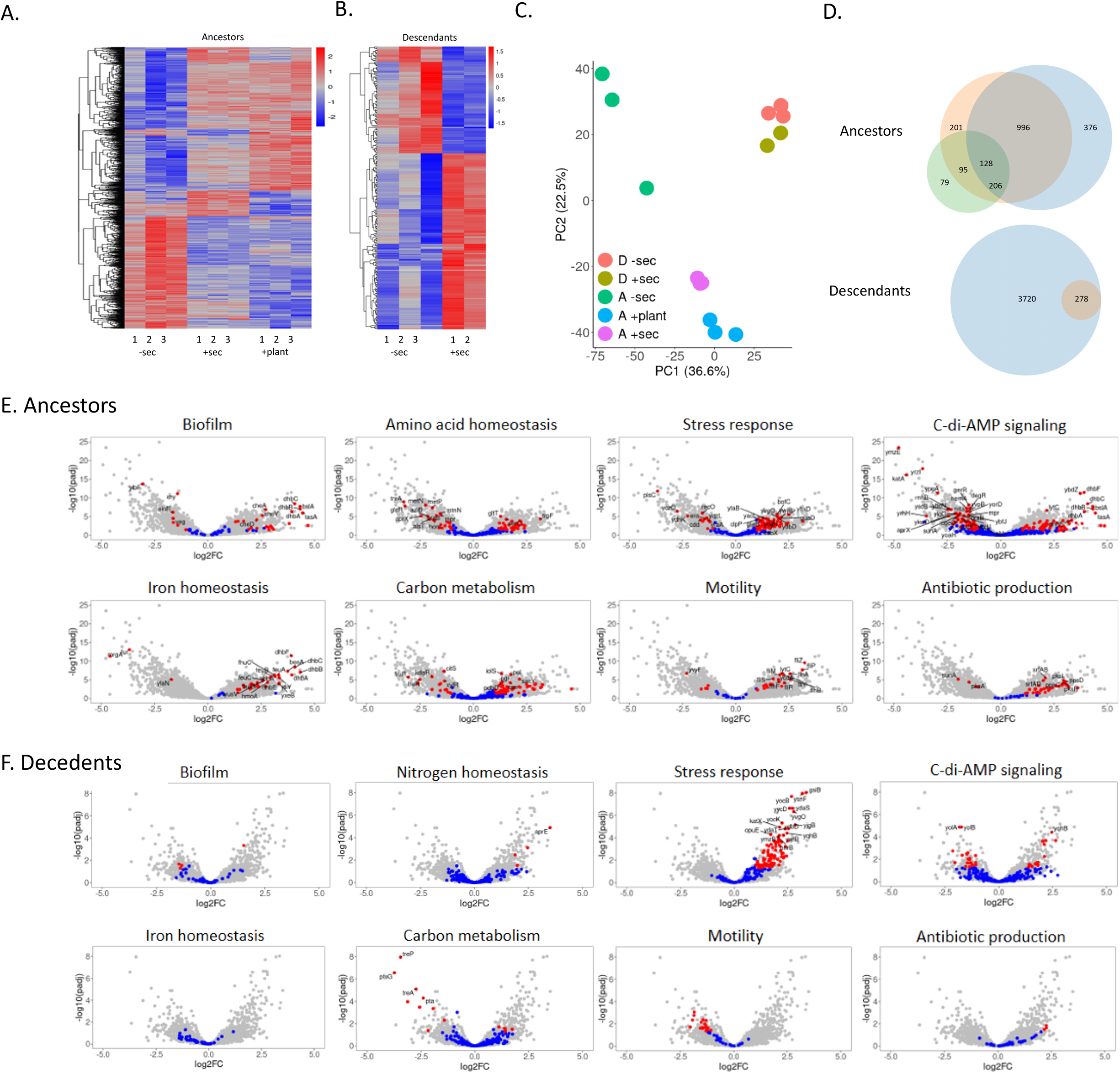
Multigenerational adaptation of *B. subtilis* to *A. thaliana* involves transcriptome rewiring Long-term adaptation of *B. subtilis* to A. thaliana involves transcriptome rewiring. (A) Hierarchical clustering of genes in ancestor and descendant bacteria that were detected as differentially expressed in the transcriptome experiment. Each treatment includes three replicates.. The data are presented on a Z-score scale. B. subtilis in the absence of secretion treatment (-sec), in the presence of secretions (+sec), and the presence of A. thaliana seedlings (+plant), as well as descendants of secretion treated and untreated bacteria. (B) Principal component analysis (PCA) of the RNA-Seq data of untreated, secretion treated, and plant treated ancestor bacteria (“A”) and their descendants (“D”). (C) Venn diagrams of differentially expressed genes in the samples mentioned above. The number in each circle represents the amount of differentially expressed genes between the different comparisons and their overlap: – sec and + sec (red circle), -sec and +plant (blue circle), +sec and +plant (green circle) in ancestors. In the descendants, the red circle represents the number of differentially expressed genes out of all the genes (blue circle). (E-F) Volcano plots. The red dots represent differentially expressed genes between secretion treatment and untreated cells, and the blue dots represent non-differentially expressed genes in the mentioned gene categories. The other genes are represented by the light gray dots.

**Figure 5.**
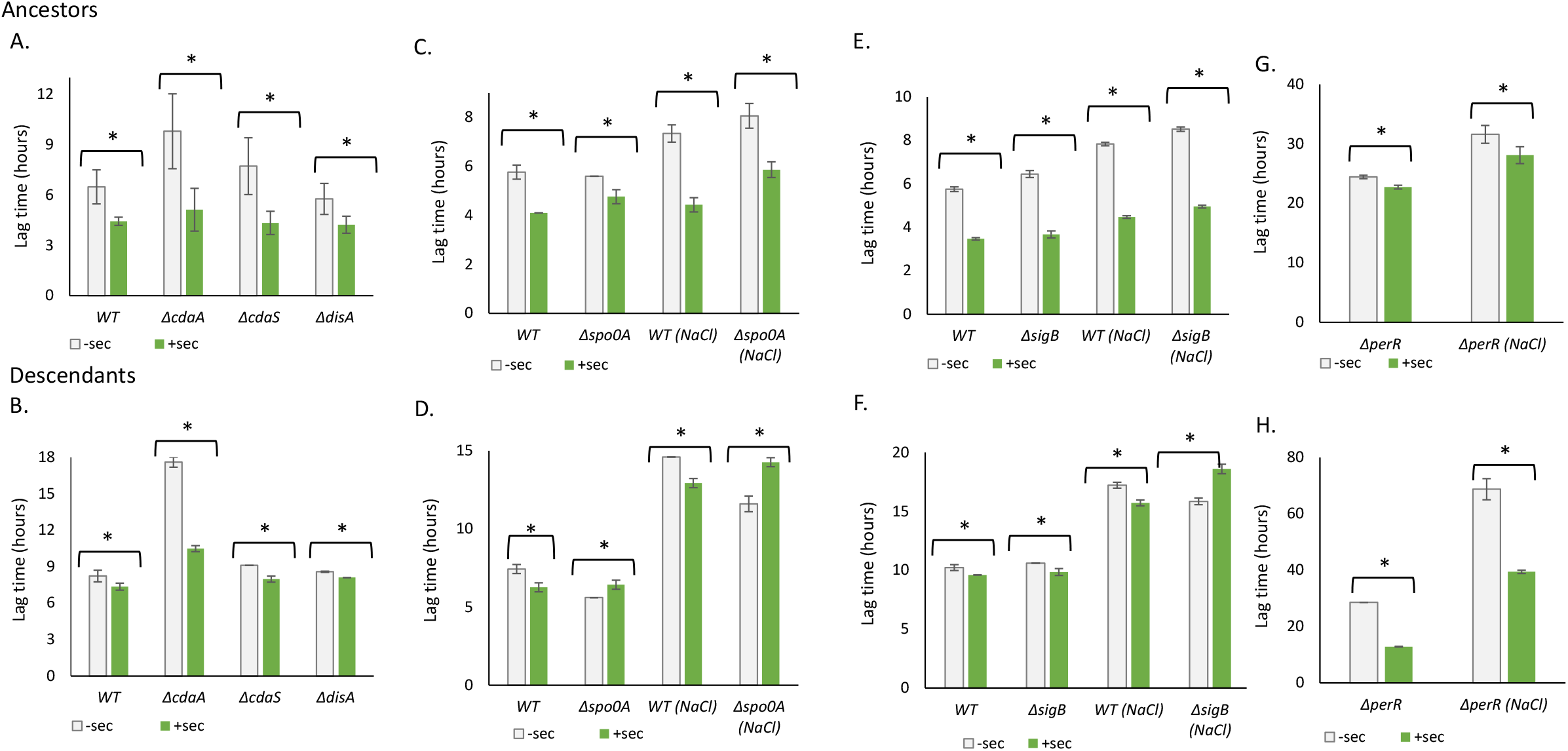
Multigenerational adaptation of *B. subtilis* to *A. thaliana* involves nucleotide signaling, the general stress response and the master regulator Spo0A_. (A-B) Lag time of WT versus diadenylate cyclase mutant bacteria. WT, *ΔcdaA, ΔcdaS* and *ΔdisA* Bacteria were treated or untreated with plant secretions (+sec and -sec respectively). The bacteria and their descendants were grown in a microplate reader and their lag time was calculated (See Fig.1). (C-H) Lag time of secretion-treated or untreated WT, Δ*sig*B, Δ*spo0A* and Δ*per*R bacteria (C-E) and their descendants (F-H). The bacteria and their descendants were grown in a microplate reader in the absence and presence of NaCl and their lag time was calculated (See Fig.1). Data is expressed as average values ± standard deviations of the means from a representative experiment, performed with triplicates. Data was analyzed by two tailed student’s t-test (alpha = 0.05).

The plant could directly provide c-di-AMP or alternatively, plant secretions may modulate the intracellular levels of c-di-AMP. Exogenously applied c-di-AMP did not mimic the effect of plant secretions, as the growth curves of the treated bacteria overlapped those of untreated bacteria (Fig. S9), raising the hypothesis that the plant may modulate the intracellular levels of c-di-AMP indirectly. In agreement with this hypothesis, the expression of the genes encoding the diadenylate cyclase DisA and the c-di-AMP transporter YhcA was induced by pre-exposure to root secretions, and *cdaS* was elevated in their descendants (supporting data sheet 1). These results are consistent with the capacity of the plant to compensate over the loss of the adenylate cyclase CdaA. In addition, we observed a differential expression of genes that are involved in potassium homeostasis, and thereby to osmotic stress tolerance such as *kimA*, encoding a potassium transporter, *yjbQ* potassium exporter and *khtT*, a K+/H+ antiporter^36^, whose activities is also modulated by direct c-di-AMP binding^37^ (Fig. S10). These results raise the hypothesis that plant alters c-di-AMP levels and thereby induces a long-term general stress response.

The stress response in *B. subtilis* can be activated by the general stress response regulators (primarily the alternative sigma factor SigB^38^) and the master regulator Spo0A^39^, both acting downstream to c-di-AMP signaling^30^. Some of the target genes of SigB overlap with the target genes of PerR, a dimeric repressor that regulates the response to peroxide stress, and thereby PerR regulon is overlapping at some extent with SigB regulon ^40^. Therefore, we evaluated the major regulators of stress tolerance by evaluating the response of their, deletion mutants to plant secretions. We first evaluated the requirements for Spo0A, as a major regulator of plant colonization^14^. Root secretions reduced the lag time of a mutant lacking Spo0A to a smaller extent compared with the WT parental strain, both in the absence and presence of NaCl, while their descendants exhibited a slight increase in lag time (e.g: an opposite response to the plant) following secretion treatment in the presence and absence of salt (Figs 5C-D). As Spo0A deletion did not eliminate the beneficial effects of plant secretions, we tested the general stress response main components: Secretion-treated Δ*sig*B strain was comparable to WT in the absence and presence of NaCl, while Δ*per*R was largely resistant to root secretions compared with the parental strain, and a *sig*B mutant (Figs 5E and G). These results indicate that plant secretions exert their specific early effects primarily through PerR. In contrast to the presence of significant response of ancestors lacking the *sig*B to secretions, decedents of the mutant did not show an increased tolerance to stress in response to secretions. Plant secretions increased (rather than decreased) the lag time of *sigB* mutant descendants of treated bacteria, indicating that SigB absence does not allow plant secretions to convey late beneficial effects in the exposed bacteria (Fig. 5F). Similarly, in contrast to the lack of response of ancestors lacking PerR, in the descendants, *per*R mutant exhibited a pronounced reaction to secretions than WT in the absence and presence of NaCl, indicating that PerR participates in sensing early but note late effects of plant secretions (Fig. 5H).

Overall, these results indicate that root secretions trigger the *B. subtilis* stress response and require PerR/SigB and Spo0A, supporting the differential expression of their regulons.

### *A. thaliana* secondary metabolites can exert beneficial multigenerational response for *B. subtilis*

To identify the plant compound(s) responsible for the phenotype we observe in the bacteria following secretion treatment, we used the parental strain bearing *PpksC* fused to a luciferase reporter. The choice of this reporter resulted from our finding that *pks* operon genes were significantly upregulated in secretion-treated bacteria and their descendants (Fig. S11). These results are consistent with our previous findings that *A. thaliana* and *Eruca sativa* induce *pks* expression ^41,42^.

We first monitored the planktonic growth and *pks* promoter expression in the presence of several plant metabolites that modulate plant-bacteria interaction, i.e., salicylic acid, which induces the SigB-dependent general stress response in *B. subtilis* ^43^ and jasmonic acid, whose derivative methyl jasmonate affected the rhizosphere community of *A. thaliana ^44^*. We also tested again sucrose, which promotes *B. subtilis* plant colonization ^15^, and malic acid, a plant derived preferred carbon sources for *B. subtilis ^45^*. None of those mentioned above compounds mimicked the effect of plant secretions on *pks* promoter expression (Fig. S12).

Finally, we screened secretions from *A. thaliana* strains with an aberrant accumulation of various secondary metabolites for both plant-derived benefits, which are sustained over time: the induction of the bacterial *pks* promoter and shortening of the lag phase. We screened mutants in genes coding for enzymes in the methylerythritol phosphate (MEP) pathway, i.e., DXR and DXS. This pathway produces isopernoids metabolites with essential functions in plants and bacteria^46,47^. We also examined secretions from an *Arabidopsis aberrant lateral root formation 4* (*alf4*) mutant that has an aberrant auxin signaling^48^ and a mutant deficient in one of the lipoxygenases (LOXs), which are involved in the biosynthesis of a large group of biologically active fatty acid metabolites collectively named oxylipins^49^.

We found that secretions from a DXS overexpressing strain (*dxs* OE) upregulated the expression of the *pks* promoter to a similar extent as WT secretions. However, secretions a DXS-deficient strain (*dxs1*), failed to replicate the effect of WT secretions on *pks* expression (Fig. 6A-C). Furthermore, secretions from all but a single mutant strain (*lox2-1*) yielded a milder impact on the bacterial lag phase than WT secretions (Fig. 6D-F).

**Figure 6.**
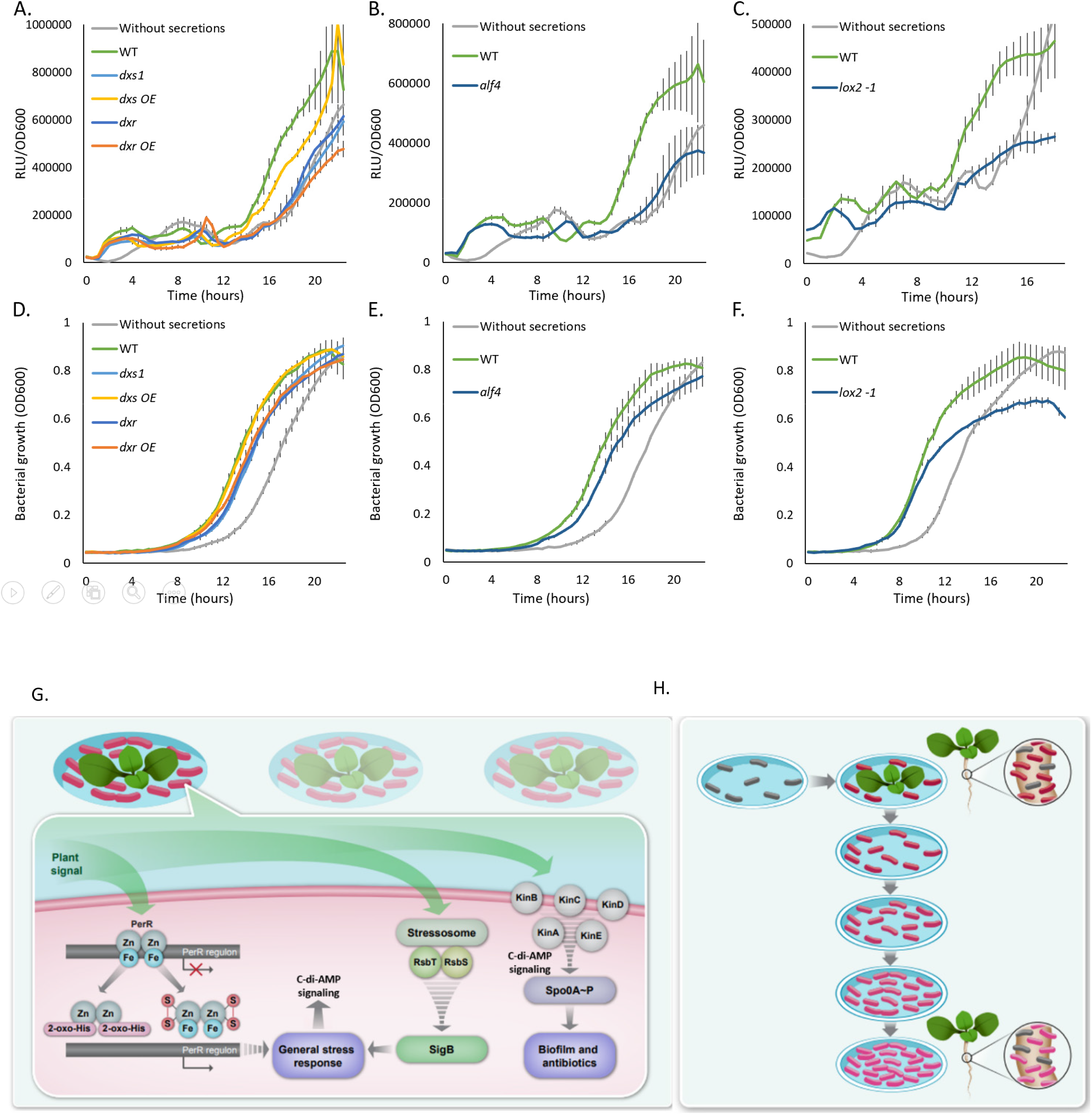
Oxylipins support the beneficial effects of A. thaliana. (A-C) Normalized expression of PpksC-lux and (D-F) growth curves of bacteria grown in the absence or presence of WT and mutant A. thaliana secretions. Data is expressed as average values ± standard deviations of the means from a representative experiment, performed with four technical repeats. Data was analyzed by two tailed student’s t-test (alpha = 0.05). **(G) A proposed bacterial pathways affected by the plant**. (i) The plant modulates the activity of Spo0A and its kinases, which regulate biofilm formation and antibiotic synthesis. (ii) The plant modulates the activity of SigB, whose activity is phosphorylation-dependent and involves the stressosome via one of several putative sensor proteins. SigB, in turn, controls the general stress response. (iii) The plant modulates the activity of PerR by one of the two scenarios indicated. PerR modulates the response to peroxide stress and indirectly affects the general stress response. **(H) A scheme showing the benefits of memorization of plant hosts.** Naïve bacteria (grey) are treated with a plant or its secretions. The treated bacteria (red) then colonize the root more efficiently than untreated bacteria in a competition setting. The effect of the plant on the bacterial transcriptome is partially conserved in the descendants of the treated bacteria (pink). However, they still exhibit a higher efficiency of root colonization, which is comparable to their ancestors.

These results support a possible role of plant isoprenoid and oxylipin biosynthesis and auxin signaling in modulating the bacterial lag phase and *pks* expression. These pathways could have a complementary function in inducing the bacterial phenotype we observe following interaction with the plant.

## Discussion

*B. subtilis* is a promising sustainable alternative to traditionally used hazardous pesticides, as it can protect crops from fungal, bacterial, and viral pathogens, without harming the diversity of their environment to the same extent. The beneficial properties of this bacterium and related species are attributed to the production of a wide arsenal of antibiotics and antimicrobial substances with a broad repertoire ^50^. Currently, while biocontrol agents belonging to the *B. subtilis* clade carry tremendous potential for agriculture the results in the field often fail to compare with chemical pesticides^51^. Therefore, to successfully applicate strains of *B. subtilis* and related specie in agriculture, the resolution of the underlying mechanisms of how it interacts with and colonizes plants is extremely useful.

Under standard laboratory settings, several cues have been associated with biofilm maturation and assembly, including oxygen deprivation ^52^, nutrient deprivation ^53–55^, small molecule sensing^30,56,57^, calcium ^58,59^ and physical cues ^60–63^. In addition, it was shown that plant polysaccharides, and malic acid may act as signals to induce biofilm formation ^15,64^ that plant secretions can enhance chemotaxis ^16^, and that sucrose from the plant can alter gene expression to promote root colonization ^17^.

It is interesting to inquire whether adaptation to the plant can also be manifested by long-term memory. In recent years, evidence emerged that bacteria contain complex control circuitry capable of generating multi-stable behaviors and other complex dynamics that have been conceptually linked to memory in other systems ^65^. It has long been known that bacterial cells that have experienced different environmental histories may respond differently to current conditions^66^. In addition, genetic diversification of *B. subtilis* into morphotypes also plays an important role in evolution during plant root colonization ^25^. Considering the complex nature of plant-bacillus interactions, history-dependent behavioral differences may be physically necessary consequences of the prior interaction, as they may stabilize symbiosis by providing ongoing benefits to the interacting bacteria.

Here, we could demonstrate such multigenerational effect from interaction with the host: First, bacteria pre-exposed to plant secretions and their descendants (grown for 8 generations in a rich media) exhibited and improved growth and enhanced stress tolerance to both salinity stress and high concentrations of the plant phytohormone salicylic acid. Second, and potentially due to this increased fitness, the exposure to the plant root increased the capacity of the ancestors and descendants to colonize the plant efficiently. Overall, this enhanced fitness was robustly manifested by the capacity of pre-exposed bacteria and their descendants to compete with naïve bacteria for the colonization of a new root.

Transcriptome analysis to an exhaustive set of culturing experiments on *B. subtilis* demonstrated that *B. subtilis* experience short term and long term changes in gene expression following its exposure. More short term than long-term memory was evident in response to the plant (Fig. 4). However, 278 genes were still differentially expressed in the descendants of pre-exposed bacteria, and many of them belonged to either the general stress response, or c-di-AMP signaling. Both pathways are possibly interlinked, as the diadenylate cyclase DisA is part of the general stress regulon of SigB ^67^. Our hypothesis that the plant metabolites actively trigger the general stress response via altering c-di-AMP signaling is supported by our finding that plant secretions can compensate for the loss of the diadenylate cyclase CdaA by inducing DisA (in ancestors) and CdaS (in descendants). Furthermore, descendants of secretion-treated Δ*sig*B bacteria do not respond to NaCl in contrast with the WT parental strain (Fig. 5), indicating that the general stress response is required to guarantee the long-term effects on descendants of bacteria that interacted with the root. Indeed, both ancestors and descendants of bacteria that interact with root secretions become more tolerant to biotic stress, represented by sialycilic acid, and abiotic stress represented by high salinity (Fig. 2). These simple experiments and the surrounding analysis and framework demonstrate what could be the beginning of a larger multigenerational transcription rewiring program, and indicate that bacterial memory in cellular behaviors may be a rich area for further exploration.

Consistent with the established role of Spo0A in plant colonization, *spo0A* deletion manifested a prolonged lag time as compared with WT bacteria and their descendants. In addition, we found an aberrant response to plant secretions in *per*R knockout (Fig. 5). Taken together, these results support the notion that plant-derived compounds act upstream of SigB and Spo0A, by modulating c-di-AMP signaling (Fig. 6G).

Admittedly, we do not know yet what plant signal(s) are responsible for the observed pattern of multigenerational information storage. Furthermore, metabolites produced by the root (e.g salicylic acid, sucrose, malic acid and jasmonic acid) that were previously reported to induce microbial responses did no mimic the long-term effects of plant secretions (Fig. S12). It is possible that a combination of several metabolites is required in order to induce the effect we observe in the bacteria following secretion treatment. Encouragingly, a mutant for oxylipins synthesis demonstrated a failure to shorten the lag phase and to induce a reporter for bacillaene synthesis, which is associated with both short-term and long-term effects of the plants (Fig. 6). As neither carbon or nitrogen mimicked the effects of the plant (Figs S2 and S4), collectively these results support a highly specific regulation by plant metabolites.

In conclusion, our results suggest that *B. subtilis* adapts rapidly to the interaction with *A. thaliana*, and that this adaptation includes unexpected multigenerational transcriptome rewiring and increased fitness. Our findings are relevant for the application of *B. subtilis* as a PGPR in agriculture, as they imply that simple pre-exposure of the inoculated strains to the root might be a stable strategy for root colonization and improved stress tolerance of the applied strains (Fig. 6H). Moreover, an increased multigenerational fitness following an exposure to a host cue could be also applicable to *Bacillus* strains acting as mammalian probiotics.

## Materials and methods

### Strains and media

All bacterial strains used in this study appear in table S1. Transformation of gene knockouts from strains 168 or PY79 into strain 3610 was done with the following protocol: DNA was extracted from strain 168 or PY79 with the DNeasy Blood & Tissue kit (Qiagen) as described in the manufacturer’s protocol and the final DNA yield was quantified by Qubit (Life Technologies). The DNA was introduced by transformation into strain 3610 by natural competence, and strains were confirmed by PCR for successful transformation. The strains were grown in LB broth (Difco), or MSgg medium (5 mM potassium phosphate, 100 mM MOPS (pH 7), 2 mM MgCl2, 50 μM MnCl2, 50 μM FeCl3, 700 μM CaCl2, 1 μM ZnCl2, 2 μM thiamine, 0.5% glycerol, 0.5% glutamate, 50 μg/mL threonine, tryptophan and phenylalanine). Solid LB medium contained 1.5% bacto agar (Difco) (Maan & Gilhar et al., 2021).

### Determination of cell density

For culture density, OD600 was measured with a spectrophotometer (Ultrospec 2100, Amersham Biosciences).

### Plant strains and growth conditions

Seeds of A. thaliana Col-0 and Lycopersicon esculentum (tomato, cultivar moneymaker) were surface-sterilized and seeded on Petri dishes containing Murashige and Skoog medium (4.4 g/L), PH 5.7, supplied with 0.5% (w/v) plant agar (Duchefa), 0.5% Sucrose (SigmaAldrich) and then stratified at 4°C for two days (Golani et al., 2013). The seeds were further transferred to a growth chamber (MRC) at 23°C ia n 12 h light/12 h dark regime (Maan & Gilhar et al., 2021). All A. thaliana mutant lines were in the Col-0 background and were kindly provided by Asaph Aharoni’s lab, Department of plant and environmental sciences, Weizmann institute of science.

### Extraction of plant secretions

10-day-old seedlings were washed in PBS (Biological Industries), transferred to 6 ml liquid MSgg medium of each well of a 6-well microplate (Thermo Scientific), and then grown for additional four days. Eight seedlings were put in each well. The liquid medium was then collected and filtered with a 0.22 μm filter.

### Bacterial culture with and without plant secretions

B. subtilis cells were grown from a single colony isolated over LB plates to a mid - logarithmic growth phase (4 hours at 37°C with shaking). The cells were washed in PBS and cultured at OD of 0.1 in a 24-well plate containing liquid MSgg (“untreated”) or plant secretions for 4 hours.

### Growth measurements

Bacterial cells, treated or untreated with plant secretions, were centrifuged and washed three times in PBS. The bacteria were then seeded at an OD of 0.01 in 200 μl liquid MSgg medium of each well of a 96-well microplate (Thermo Scientific) in the presence or absence of 600 mM NaCl. The cells were grown with agitation at 30°C for 30 hours in a microplate reader (Infinite M Plex; Tecan, Austria, or Synergy 2; BioTek, Winooski, VT, USA), and the optical density at 600 nm (OD600) was measured every 30 min.

### Luminescence experiments

Luminescence reporter was grown in either MSgg medium or MSgg medium containing plant secretions. Experiments were carried out using a flat bottom 96-well plate with opaque white walls (Corning). Measurements were performed every 30 min at 30°C for 30 h, using a microplate reader (Infinite M Plex; Tecan, Austria, or Synergy 2; BioTek, Winooski, VT, USA). Luciferase activity was normalized to avoid artifacts related to differential cell numbers as RLU/OD^42^

### Lag time calculations

The bacterial lag time was calculated with the software GrowthRates 3.0 ^68^

### Bacterial root colonization measurements

Bacterial cells either treated or untreated with plant secretions were centrifuged, and cell pellets were washed three times in PBS. The bacteria were then seeded at an OD of 0.02 in 1 ml liquid MSgg with 7-d-old A. thaliana seedlings in a 12-well plate (Thermo Scientific) or in 2 ml liquid MSgg with 7-d-old tomato seedlings in a 6-well plate (Thermo Scientific). Each well contained one seedling. After 24h culture, the seedlings were removed from the liquid medium and washed in PBS. The roots were removed, transferred to a 1.5 Eppendorf tube in 1 ml of 10mM MgCl2, and vortexed for 1 minute. The bacteria were further detached with a sonicator (BRANSON digital sonifier, Model 250, Microtip) at an amplitude of 10%, pulse 3x 10 seconds.

### Colony forming units’ analysis

To determine the number of live cell counts, cells were serially diluted in PBS, plated on LB plates, and colony □forming units (CFU) were counted after incubation at 30°C overnight.

### Flow cytometry

Cells were harvested from plant roots and sonicated using the procedure discussed above. For flow cytometry analysis, cells were suspended in PBS and measured on a BD LSR II flow cytometer (BD Biosciences). The GFP and mKate fluorescence were measured using laser excitation of 488 nm and 561 nm (respectively), coupled with 505 LP and 525/50 sequential filters. The photomultiplier voltage was set to 484 V. 100,000 cells were counted, and every sample was analyzed with Diva 8 software (BD Biosciences).

### Confocal microscopy

Plants were cultured with secretion treated and untreated bacteria for 24 hours as described in the “bacterial root colonization measurements” section above. The plant roots were then washed in PBS and mounted on a microscope slide, and covered with a poly-L-Lysine 31 (Sigma)-treated coverslip. Cells were visualized and photographed using a laser scanning confocal microscope (Zeiss LSM 780) equipped with a high-resolution microscopy Axiocam camera. Data were captured using Zen black software (Zeiss, Oberkochenm, Germany) (Maan & Gilhar et al., 2021).

### RNA extraction

For RNA extraction, cells were treated with 250 μl 20g/L lysozyme for 20 minutes at 30oC. RNA extraction was done with TRIzolTM reagent (InvitrogenTM according to the manufacturer’s protocol. Briefly, 1 ml of TRIzol reagent was added, and cells were incubated at room temperature until phase separation. Then 0.2 ml of chloroform was added. The sample was centrifuged for 15 minutes at 12,000 RCF at 4oC. The aqueous phase containing the RNA was transferred to a new tube. 0.5 ml of isopropanol was added, and the samples were incubated for 10 minutes. Samples were centrifuged for 10 minutes at 12,000 RCF g at 4°C and the pellet was kept and washed in 1 ml of 75% ethanol. The samples were vortexed briefly and then centrifuged for 5 minutes at 7500 RCF at 4°C. The supernatant was discarded, and sample quality was analyzed by AgilentTM TapestationT M and NanoDropTM 2000.

### DNase treatment

TURBO DNA-free (Ambion) kit was used to remove DNA contamination, as manufacturer’s protocol. Total RNA (5μg) of each sample was digested with 1 μl DNase at 37oC for 30 minutes. Next, 0.1 of the total volume of DNase Inactivation Reagent was added, and samples were incubated for 5 minutes at room temperature. The samples were centrifuged at 10,000 RCF for 1.5 minute,s and the supernatant was kept at 4°C.

### rRNA depletion

Ribo-Zero rRNA Removal bacterial Kit (Illumina) was used to deplete the ribosomal rRNA from all samples as described in the manufacturer’s protocol. The magnetic beads were washed as described in the protocol. The probes were hybridized into the sample, and later removed with the magnetic beads according to the protocol. Ethanol precipitation was used to clean up the depleted RNA. After the process, the sample RNA concentration was measured with Qubit® 3.0 Fluorimeter.

### Transcriptome sequencing

The preparation of the libraries and sequencing were performed by the G-INCPM unit at the Weizmann Institute of Science. Libraries were prepared with G-INCPM mRNA-Seq protocol without polyA capture. Sequencing was performed by HISeq Illumina Sequencer using high output mode. Read type was NextSeq 75 cycles single pair with 200 x 106 reads. Since we had 15 samples in the lane, we expected to have around 1 x 106 reads per sample.

### RNA-Seq data analysis

Fastq reads were trimmed from their adapter with cutadapt and aligned to the B. subtilis genome (subsp. subtilis str. NCIB 3610, NZ_CM000488.1) with Bowtie 2 version 2.3.4.1 (1). The number of uniquely mapped reads per gene was calculated with HTSeq (2). Normalization and testing for differential expression were performed with DESeq2 version 1.16. Genes with normalized mean read ≥ 50, fold change ≥ 2, and adjusted P < 0.05 were considered as differentially expressed. DAVID (Database for Annotation, Visualization, and Integrated Discovery) (3-4) was used to perform gene annotation enrichment analysis.

### Statistical methods

All studies were performed in technical triplicates with three separate and independent times. Statistic analysis is provided in the legend of each figures. Data is expressed as average values ± standard deviations of the means. Data was analyzed by two tailed student’s t-test (alpha = 0.05) unless indicated otherwise.

