## Supporting figures and table for "*Arabidopsis thaliana* induces multigenerational stress tolerance against biotic and abiotic stressors and memorization of host colonization in *Bacillus subtilis*"

### Supporting information

Supporting figures (S1-S12)

Supporting Table (S1)

Supporting references

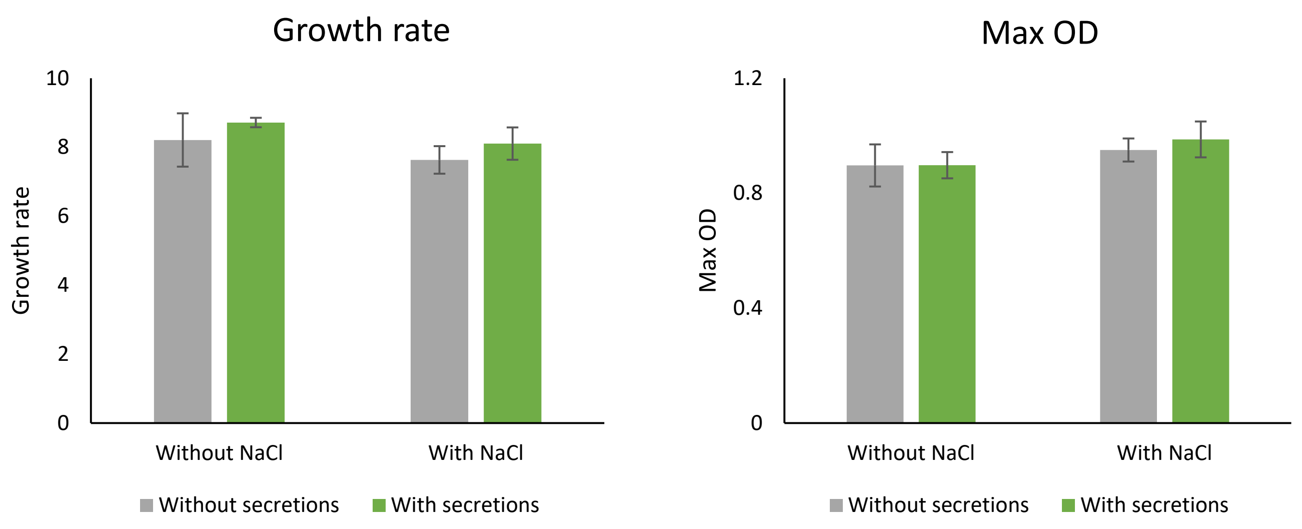


**Figure S1.** Maximal OD and growth rate of *B. subtilis* cells grown with and without pre-exposure to plant secretions, in the absence and presence of 600 mM NaCl. Bacteria were grown and treated as described in Fig. 1 and the growth rate and maximal OD were calculated (see *Materials and Methods*).

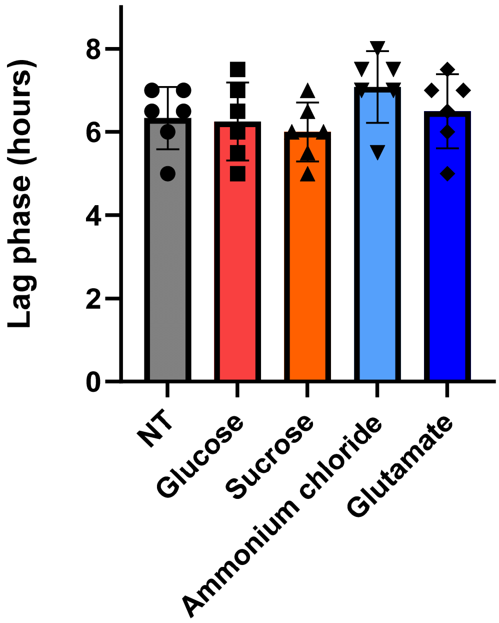


**Figure S2.** The length of the lag phase of *B. subtilis* cells with and without the indicated treatments as follows: glucose (0.1%), sucrose (0.1%), Ammonium chloride (5mM), glutamate (5mM). Bacteria were grown and treated as described in Fig. 1B and the growth rate and maximal OD were calculated (see *Materials and Methods*).

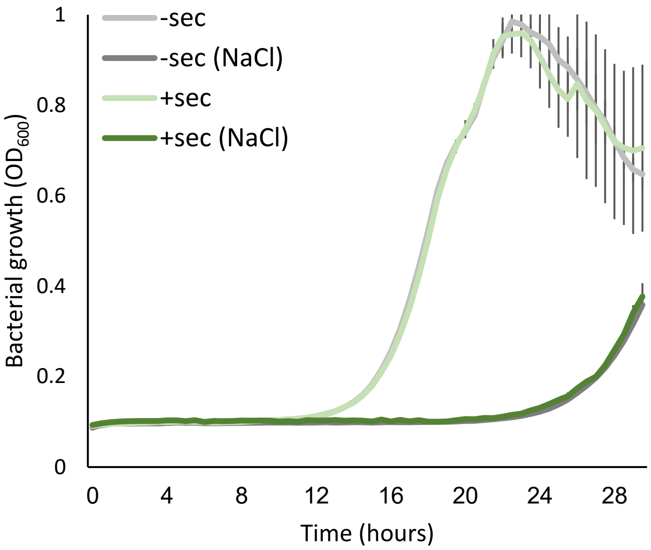


**Figure S3.** Growth curves of bacteria that were treated or untreated with secretions (+sec and –sec respectively). Treated cultures were then transferred to liquid shaking culture for 16h at $37^{o}$C and then grown in a microplate reader as described in Fig. 1.

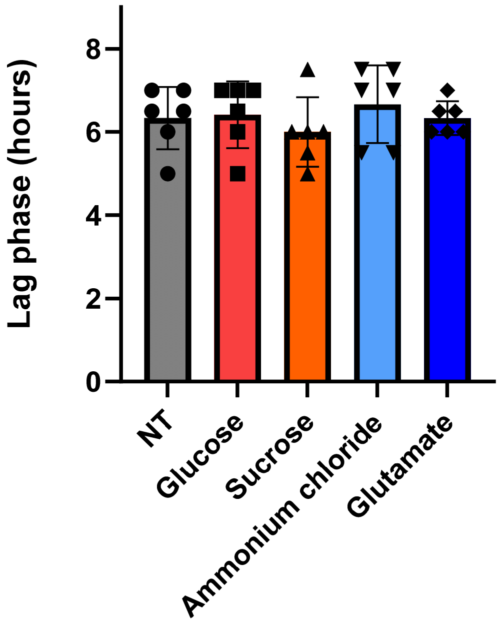


**Figure S4.** The length of the lag phase of decedents of *B. subtilis* cells originally grown with and without the indicated treatments as follows: glucose (0.1%), sucrose (0.1%), Ammonium chloride (5mM), glutamate (5mM). Bacteria were grown and treated as described in Fig. 1C and the growth rate and maximal OD were calculated (see *Materials and Methods*).

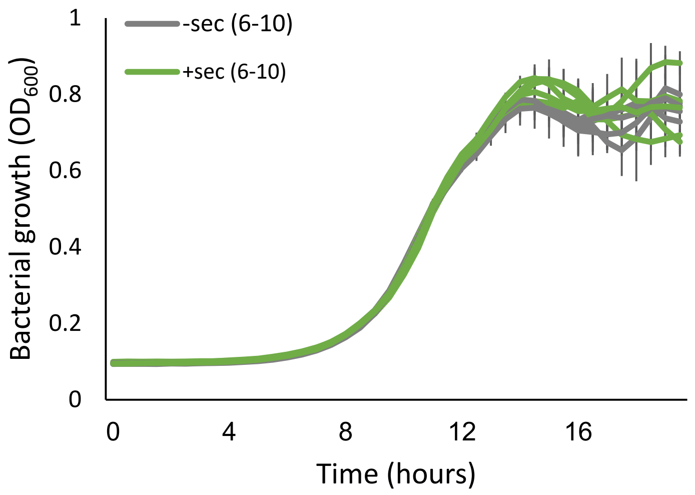


**Figure S5.** Growth curves of representative microbial colonies that were treated or untreated with secretions (+sec and –sec respectively), re-streaked separately on LB plates and grown overnight. Single colonies were picked up, transferred to liquid shaking culture for 4h at 37$^{o}$C and then grown in a microplate reader as described in Fig. 1.

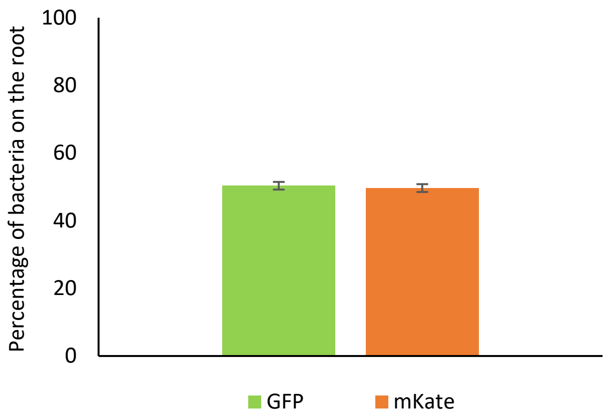


**Figure S6.** GFP and mKate labeled bacteria were cultured together in liquid MSgg with *A. thaliana* seedlings for 24h. The ratio of GFP-labeled versus mKate-labeled bacteria on the root was quantified with FACS.

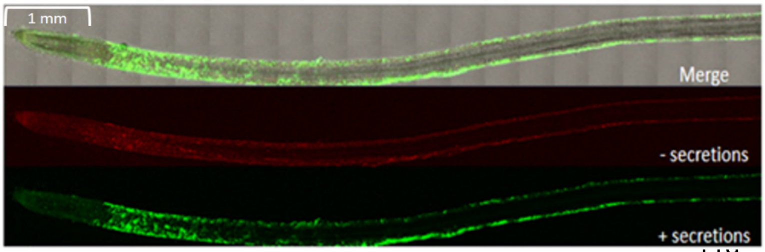


**Figure S7.** Representative bright field and fluorescent images of secretion-treated and untreated ancestor bacteria on tomato roots.

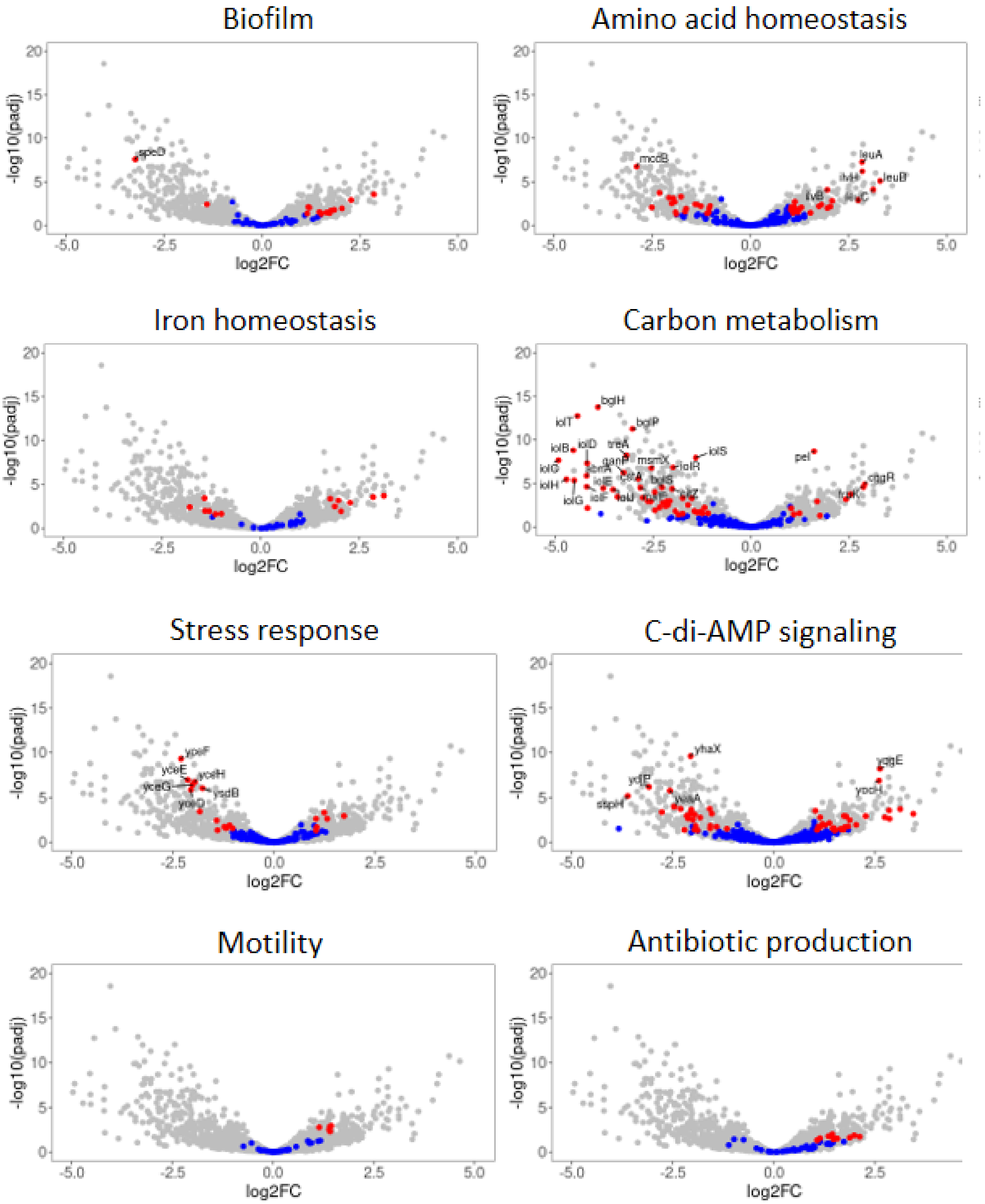
Biofilm
Amino acid homeostasis
Iron homeostasis
Carbon metabolism
Stress response
C-di-AMP signaling
Motility
Antibiotic production
-log10(padj)
log2FC
leuA
leuB
ilvB
bglH
bglP
pel
ctaA
yhaX
yqgE
ypeF
yceE
yceG
ycdB
stpH

**Figure S8.** Volcano plots of differential gene expression in bacteria cultured in the presence of *A. thaliana* seedlings as compared to bacteria cultured in secretions only. The red dots represent differentially expressed genes and the blue dots represent non-differentially expressed genes in the mentioned gene categories.

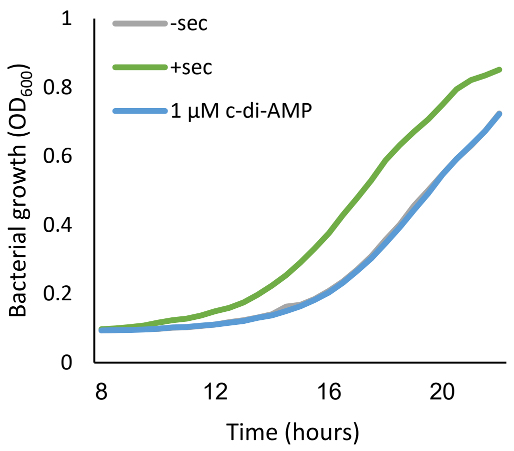


**Figure S9.** Bacterial growth in the absence and presence of c-di-AMP. Bacteria were cultured for 4h in the presence or absence of secretions (+sec and –sec respectively) and c-di-AMP and then grown in microplate reader as described in Fig. 1B.

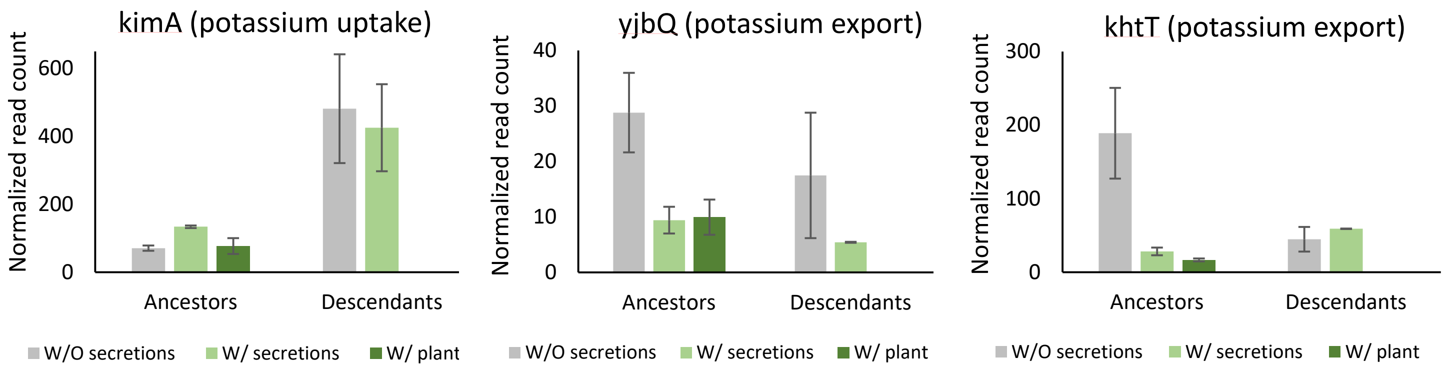


**Figure S10.** Expression estimates of genes whose expression is known to be regulated by c-di-AMP.

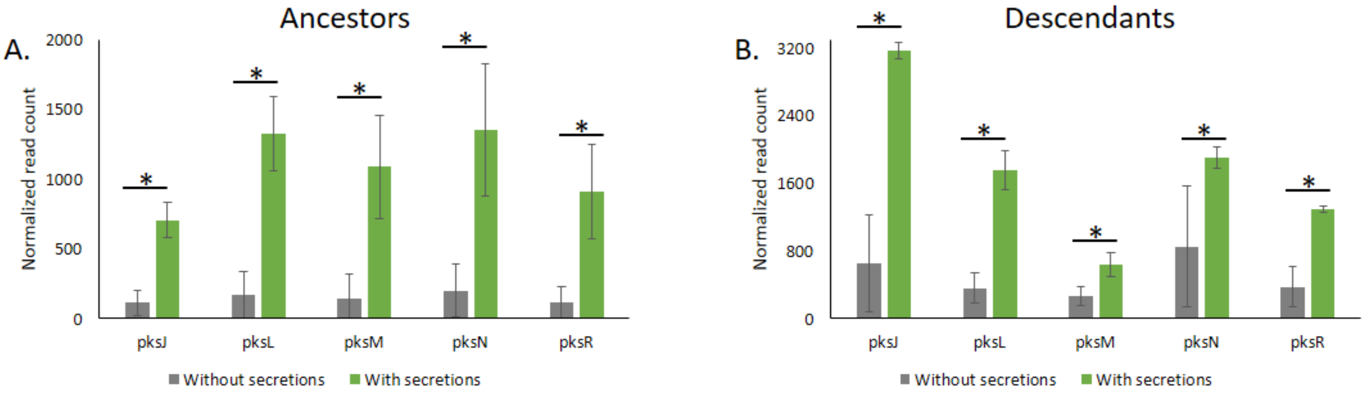


**Figure S11.** Normalized read count of pks operon genes encoding the polyketide synthetases (PKS) that synthesize Bacillaene, in the absence or presence or secretion treatment, in (A) ancestor and (B) descendant bacteria. Data were obtained from our transcriptome analysis.

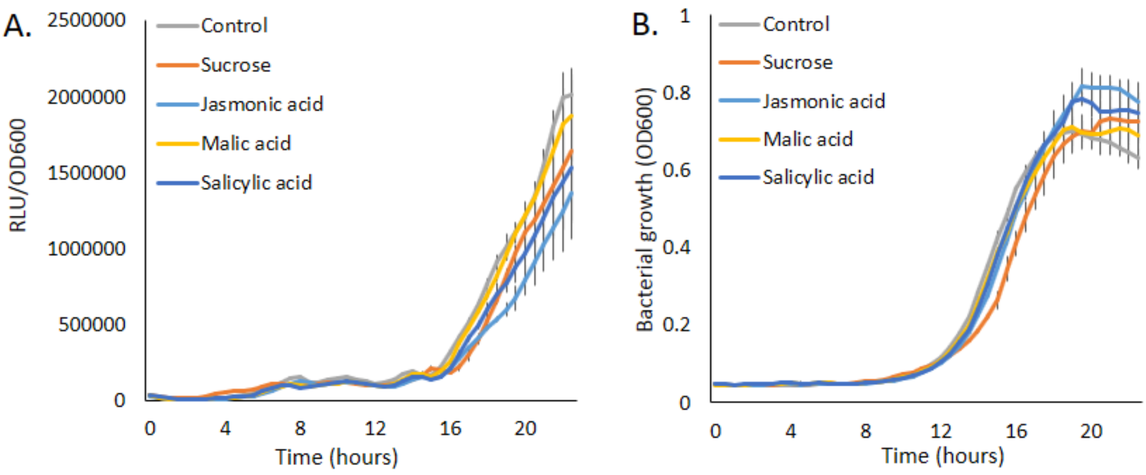


**Figure S12.** (A) The expression of $P_{pksC}$-*lux* (AU) and (B) growth of bacteria ($OD_{600}$) in the absence or presence of 100 µM of indicated metabolites.

**Table S1**

**List of strains used in this study**

| Strain | Description | Source of reference |
| --- | --- | --- |
| *B. subtilis* NCIB 3610 | Wild type | [1] |
| Δ*disA* | DNA was extracted from *B. subtilis* PY79 Δ*disA*, and transferred into *B. subtilis* 3610 (Tet$^{r}$) | Sigal Ben-Yehuda lab, Hebrew University of Jerusalem, Israel |
| Δ*cdaA* | DNA was extracted from *B. subtilis* 168 Δ*cdaA*, and transferred into *B. subtilis* 3610 (Kan$^{r)}$ | [2] |
| Δ*cdaS* | DNA was extracted from *B. subtilis* 168 Δ*cdaS*, and transferred into *B. subtilis* 3610 (Kan$^{r)}$ | [2] |
| $P_{hyperspank}$-*GFP* | Cam$^{r}$ | Lab stock, [3] |
| $P_{hyperspank}$-*GFP*-mKate | Cam$^{r}$ | Lab stock |
| Δ*sigB* | Kan$^{r}$ | Lab stock |
| Δ*spo0A* | Kan$^{r}$ | [1] |
| Δ*per*R | Kan$^{r}$ | Lab stock |
| $P_{pksC}$-*lux* | *B. subtilis* sacA::$P_{pksC}$-*lux* (Cm$^{r}$) tagged to the luciferase reporter integrated in the neutral *SacA* locus | [4] |

**Cam$^{r}$** – chloramphenicol resistance **Sp$^{r}$** – spectinomycin resistance

**Tet$^{r}$** – tetracycline resistance **Kan$^{r}$** – kanamycin resistance
